# Computing Osmotic Permeabilities of Aquaporins AQP4, AQP5, and GlpF from Near-Equilibrium Simulations

**DOI:** 10.1101/102343

**Authors:** Thierry O Wambo, Roberto A Rodriguez, Liao Y Chen

**Author notes:** Correspondence to: Liao Y Chen, Department of Physics, University of Texas at San Antonio, One UTSA Circle, San Antonio, Texas 78249 USA.

## Abstract

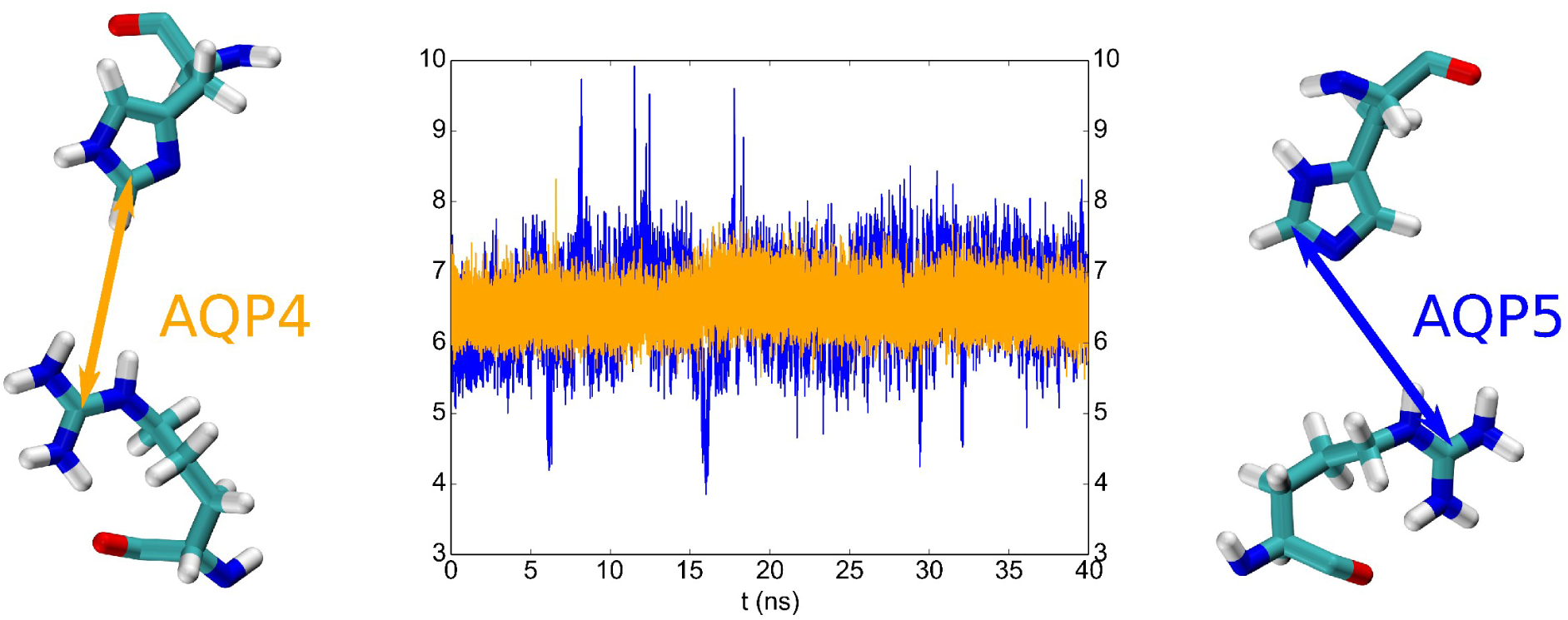

Measuring or computing the single-channel permeability of aquaporins/aquaglyceroporins (AQPs) has long been a challenge. The measured values scatter over an order of magnitude but the corresponding Arrhenius activation energies converge in the current literature. Osmotic flux through an AQP was simulated as water current forced through the channel by kilobar hydraulic pressure or theoretically approximated as single-file diffusion. In this paper, we report large scale simulations of osmotic current under sub M gradient through three water channels (the water-specific AQP4 and AQP5 along with aquaglyceroporin GlpF) using the mature particle mesh Ewald technique (PME) for which the established force fields have been optimized with known accuracy. These simulations were implemented with hybrid periodic boundary conditions devised to avoid the artifactitious mixing across the membrane in a regular PME simulation. The computed single-channel permeabilities at 5°C and 25°C are in agreement with recently refined experiments on GlpF. The Arrhenius activation energies extracted from our simulations for all the three AQPs agree with the *in vitro* measurements. The single-file diffusion approximations from our large-scale simulations are consistent with the current literature on smaller systems. From these unambiguous agreements among the in *vitro* and *in silico* studies, we observe the quantitative accuracy of the all-atom force fields of the current literature for water-channel biology. We also observe that AQP4, that is particularly rich in the central nervous system, is more efficient in water conduction and more temperature-sensitive than other water-specific channel proteins.

## INTRODUCTION

Aquaporins/aquaglyceroporins (AQPs)[1-16], the water/glycerol channel proteins, are fundamental and ubiquitous in living organisms. Naturally, these membrane proteins have been investigated in a great many experimental and theoretical/computational studies, *e.g.,* Refs. [4, 16-34] on GlpF and AQPZ expressed in *E. coli,* Refs. [8, 35-77] on AQP4, and Refs. [38, 70, 78-86] on AQP5. An essential task in the computational studies of aquaporins is to compute the channel permeability as the ratio between the osmotic current and the osmolyte concentration gradient that induces the water flux through the AQP channel, in direct parallel to the experimental measurements. Due to technical difficulties in the numerical implementation of osmotic flux induced by a sub M concentration gradient, a few studies[87, 88] have been accomplished to substitutionally compute the permeability from the water flux induced by kilobar hydraulic pressure. Due to the large pressure gradient across the membrane, these studies leave open the question about far out-of-equilibrium effects that are absent under physiological conditions. Many computational studies (e.g. Refs. [17, 22, 50, 51, 57, 79, 89-91]) have been conducted on the basis of the theoretical approximation of single-file diffusion[88, 91, 92]. The accuracy of these single-file diffusion approximations has not been ascertained because the in *vitro* measurements of single-channel permeability have proven to be very challenging as well. In fact, the experimental data on a given AQP scatter over the range of an order of magnitude. Recent experimental investigations have given us some converging data on some AQPs including GlpF[16] and AQP4[37]. In contrast to the scatter of the absolute values of permeabilities, certainty has been consistent in the *in vitro* measurements of the Arrhenius activation barrier which is rather independent of the membrane-protein expression levels and of the temperatures of the experiments. AQP1, AQP5, AQPZ and GlpF (if not inhibited by glycerol[17]) were all measured to have an Arrhenius activation barrier around 3 kcal/mol. Interestingly, AQP4 was measured to have about 5 kcal/mol, indicating this particular water channel is far more temperature-sensitive than others. (Namely, AQP4’s permeability increases more than other aquaporins when the temperature is elevated.) However, theoretical/computational studies predicted <3 kcal/mol for all cases including AQP4 in the current literature.

In this paper, we present a computational study of two water-specific channels (AQP4 and AQP5) and one water-glycerol channel (GlpF) at two temperatures (5°C and 25°C) in direct correspondence to the *in vitro* experiments. We conducted large-scale simulations of all-atom model systems in which the signal-to-noise ratios were sufficiently high to achieve unambiguous accuracy. We computed the osmotic permeability under near-physiological conditions directly as the osmotic water-current divided by the osmolyte-concentration gradient, achieving close agreement with the latest refined experimental data. We also conducted single-file diffusion approximations in all six cases (three AQPs at two temperatures) which are consistent with the current literature on smaller systems, indicating single-file diffusion approximations are not quantitatively accurate. From the temperature-dependence of the permeabilities, we extracted the Arrhenius activation barriers for the three AQPs that are all in excellent agreement with the *in vitro* results. From the differentiation between the dynamic characteristics of AQP4 and AQP5, we gained atomistic insights (structures and fluctuations) about why AQP4 is preferable for maintaining hydrohomeostasis of the central nervous system.

## METHODS

Our main objective is to achieve clear signal-to-noise ratios in direct computations of the water flux induced by a sub M osmolyte concentration gradient across the membrane. To achieve this goal, we build model systems four times as large as those in the current literature. Each model system consists of four tetramers (16 water channels) of either of the three proteins embedded in a large patch of lipid bilayer representing the cellular membrane. We compute the osmotic permeability under near-physiological conditions directly as the osmotic water-current divided by the osmolyte-concentration gradient. We employ a hybrid periodic boundary conditions (PBC) for a particle mesh Ewald (PME) implementation with the CHARMM force field.[93] In the hybrid PBC scheme, images of the system volume (cell) are arranged periodically in all three dimensions as in a usual PBC setup but a rigid plane/wall parallel to the membrane is placed at the top/bottom of the system volume to eliminate the artifactitious mixing between the aqueous volumes on the two sides of the membrane that is intrinsic in the usual PBC scheme. In this manner, we are able to maintain, for a sufficient period of time, a constant osmotic flux through 16 aquaporin channels induced by a sub M concentration gradient across the membrane. In fact, during the simulation time of 40 ns, the osmolyte concentration gradient decreased by less than 0.5% in all cases.

Shown in Fig. 1 is the all-atom model system of GlpF (PDB code: 1FX8[30]) which consists of four GlpF tetramers (16 individual monomers/channels) embedded in a patch of phosphatidylethanolamine (POPE C16:0C18:1) lipid bilayer with a layer of saline of higher concentration on the top side (z>0) and a layer of saline of 150 mM on the bottom side (z<0). The Cartesian coordinates are set up so that the xy-plane is parallel to the lipid bilayer. The usual PBCs are implemented in all three dimensions but the interfaces parallel to the xy-plane between the system and its images along the z-direction are impenetrable preventing the artificial mixing between the two sides of the lipid bilayer. An atom approaching the top and the bottom interfaces will be elastically reflected, namely, the z-component of its velocity will be inverted but the xy-components will remain unaltered. The images along the xy-directions are treated in the usual way of PBC.

**Fig. 1.**
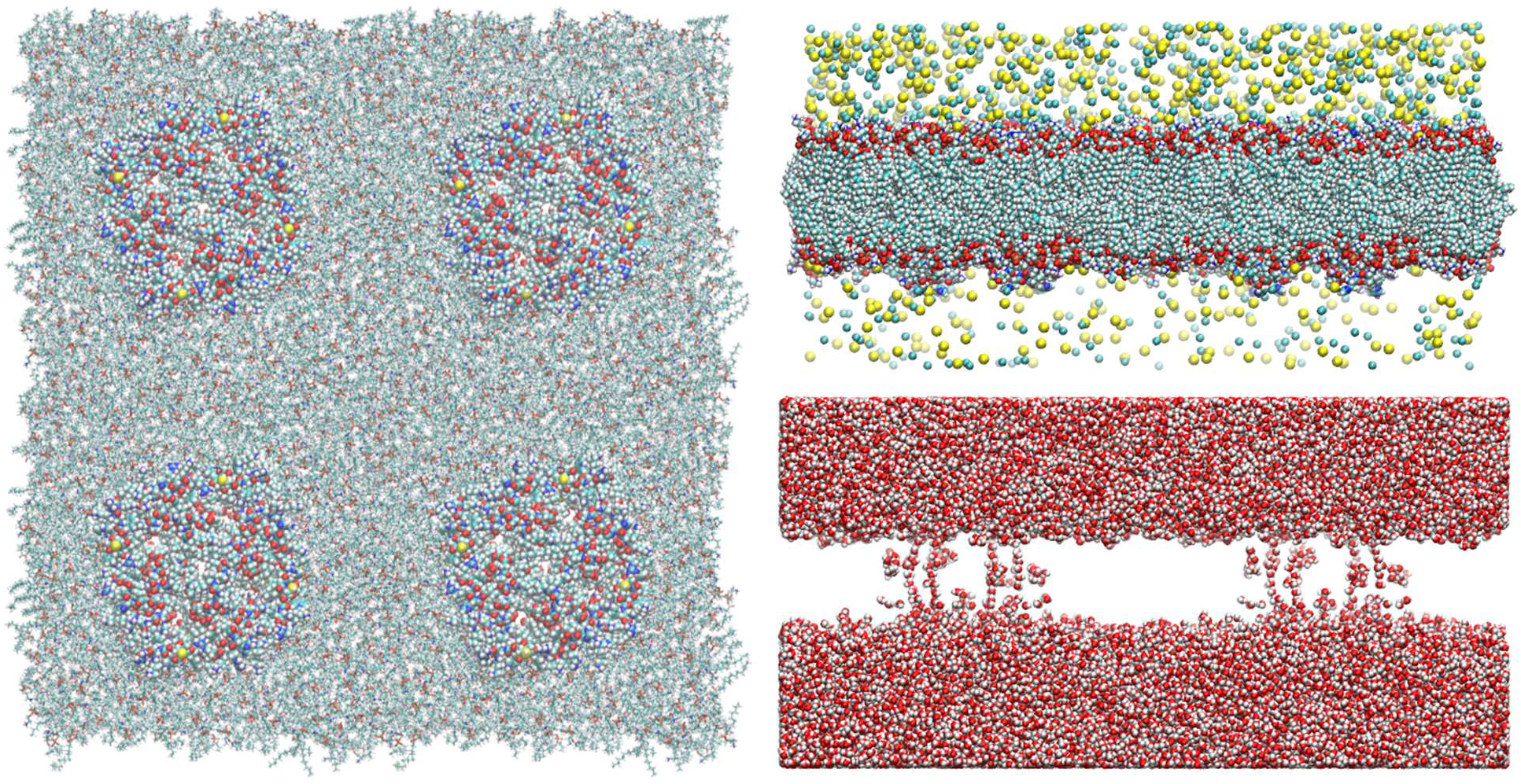
Model system of GlpF at 5°C. Shown in the left panel are the top view of four GlpF tetramers (16 individual water channels/monomers) embedded in a patch of POPE lipid bilayer. The proteins are shown as spheres and the lipids as licorices, all colored by atom names. Shown in the right panels are the side views of the GlpF model system. Top, lipids, proteins, and ions (Na+ and Cl^−^) are all shown as spheres colored by atom names. Bottom, waters are shown as spheres colored by atom names. Colors by atom names: C, cyan; N, blue; H, white; O, red; Na+, light yellow; Cl^−^, green; S, yellow; P, red. The fully equilibrated system of GlpF has the dimensions of 228*Å* × 229*Å* × 112*Å*. It consists of 601548 atoms. This and other model systems were built and the molecular graphics were rendered with VMD[94].

In order to lower computing costs, we first built a small patch (1/4 of the membrane area of the large system) with one tetramer embedded in the lipid bilayer which is sandwiched between two layers of 150 mM saline. We equilibrated the system for 100 ns with equilibrium molecular dynamics (MD) under constant temperature and constant pressure. We replicated the fully equilibrated system thrice in the xy-plane and patched them together to form the large system that has four times the membrane area of the small system. Then we added additional NaCl to the top side of the system to established an osmolyte concentration gradient across the membrane. In similar manners, we built up the model systems of AQP4-M1 (PDB code: 3GD8[8]) and AQP5 (PDB code: 3D9S[82]), which are illustrated in Figs. S1 and S2 of the supplemental information (SI). Conducting 40 ns MD runs for each of the six systems (three systems at two different temperatures), we counted the number of waters on the top side (z>0) as a function of time and used linear regression to extract the osmotic flux across the membrane through the aquaporin water pores. For the purpose of validation, we counted separately the number of waters that crossed the membrane from the bottom side to the top side but not through the 16 water channels, which amounted to less than 1% in all cases.

In all the equilibrium and nonequilibrium MD runs, we used CHARMM36[93] force field for all the intra- and inter-molecular interactions. We implemented the Langevin stochastic dynamics with NAMD[95] to simulate the systems at constant temperature of 278/298 K and constant pressure of 1 bar for equilibrium MD runs or constant volume for nonequilibrium MD runs. Full electrostatics were implemented by means of PME at a resolution of 256 × 256 × 128. The time step was 1 fs for the short-range and 2 fs for the long-range interactions. The PME was updated every 4 fs. The damping constant was 5/ps. Explicit solvent TIP3P model was used.

## RESULTS AND DISCUSSION

In Fig. 2 and SI, Figs. S3 to S5, we plot the osmotic currents of water through GlpF, AQP4, and AQP5 at 5°C and 25°C in response to a sub M osmolyte concentration gradient across the membrane. Linear regressions in all six cases give, with high confidence of fitting, the results of single-channel permeabilities which are tabulated in Table I along with relevant data from the literature. Tabulated in Table II are the Arrhenius activation barriers (energies) that indicate the temperature-sensitivities of AQP4, AQP5, and GlpF, which are in clear agreement with the experimental data of the current literature.

**Fig. 2.**
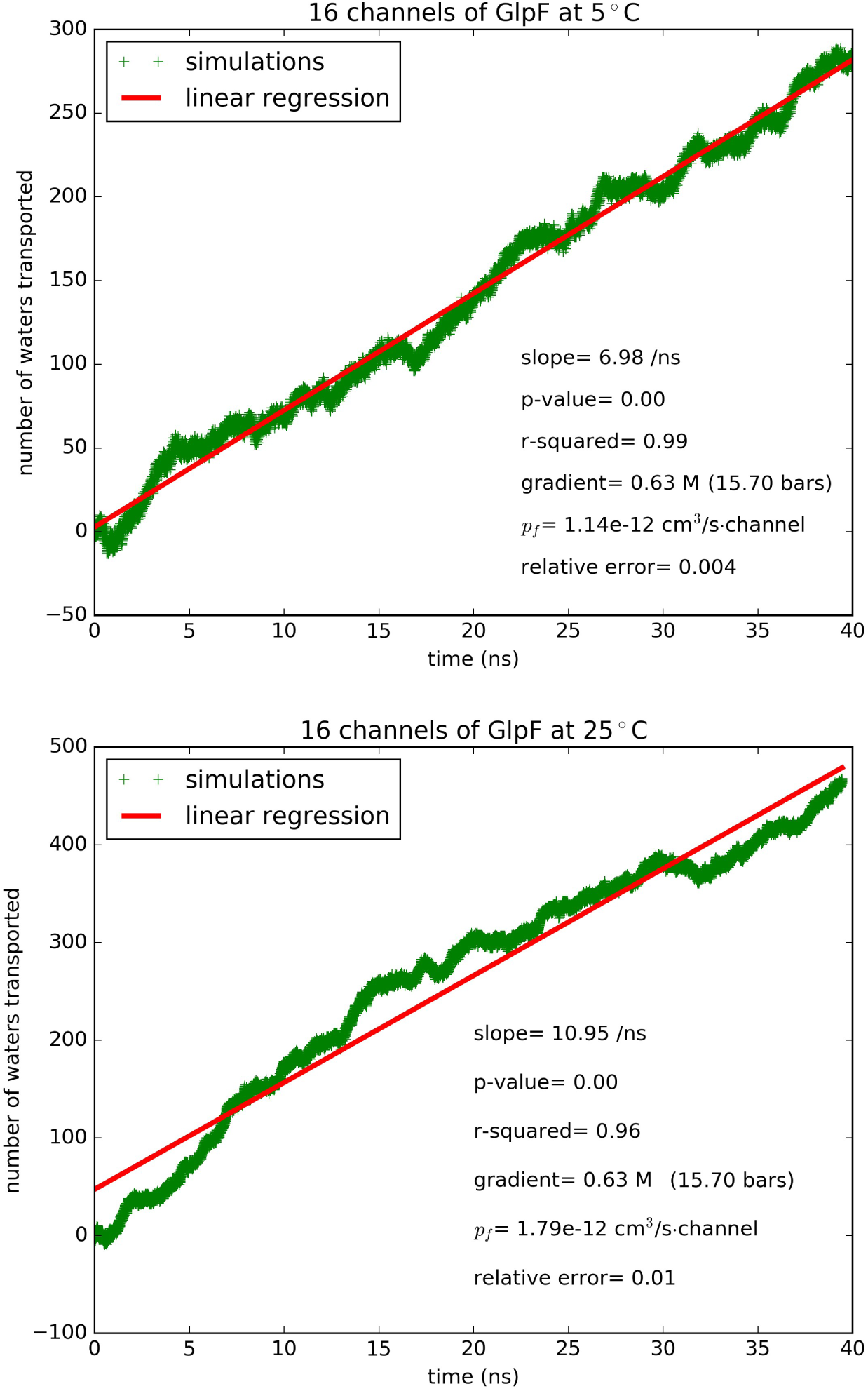
Water flux through GlpF induced by osmotic gradient at two temperatures: (A) Top panel, 5°C. (B) Bottom panel, 25°C.

**Table I.**
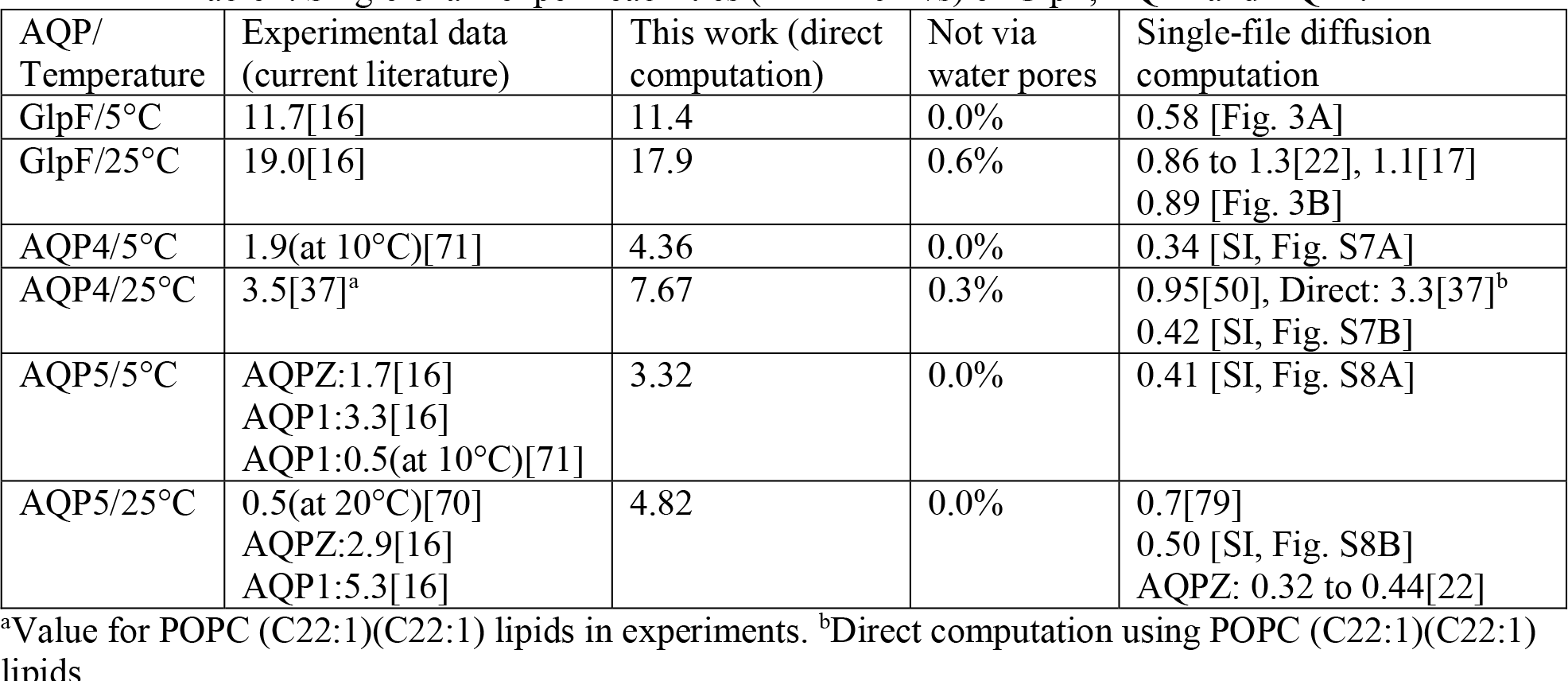
Single-channel permeabilities (in 10^−13^cm^3^/s) of GlpF, AQP4 and AQP5.

Analyzing our results in light of the current literature (Tables I and II), we observe the following three points:

### First, the single-channel permeabilities of GlpF and AQP4

The experimental data on the absolute values of the aquaporin permeabilities scatter over a range from a fraction of to more than ten times of 10^−13^cm^3^/s per single channel. The difficulty to ascertain these values has long been known for the fact that it is difficult to determine the aquaporin densities on the membrane. In a recent experimental research[16], extensive efforts led to high confidence in the extracted values of GlpF permeability. The permeability of GlpF at 5°C was found to be 11.7×10^−13^cm^3^/s per single channel[16]. Interestingly, GlpF, a glycerol channel, was thought to poorly conduct water[29]. Later on, the glycerol-GlpF interaction was computed and glycerol was identified as an inhibitor of water permeation through GlpF[17]. Now the carefully refined experiments[16] show that GlpF’s permeability is in fact very high. Our direct computation gave 11.4×10^−13^cm^3^/s (Fig. 2 and Table I), which is in perfect agreement with the *in vitro* data (11.7×10^−13^cm^3^/s).

**Table II.** Arrhenius activation barriers (kcal/mol).

| AQP | Experimental data<br>(representative values) | This work (direct<br>computation) | Computed Gibbs free-energy<br>barrier |
| --- | --- | --- | --- |
| GlpF | 4.0[16] | 3.8 | 1.5[4]; 1.7[89]; 3.4[19]; |
| AQP4 | 4.5[71];5.5[67] | 4.7 | 2.6 [SI, Fig. S9] |
| AQP5 | 3.4[80]; AQP1: 3.1[96]; AQPZ: 3[29] | 3.1 | 2.8[79]; AQP1: 1.4[4]; 3.1[19] |

Another recent experimental study[37] was on AQP4 in POPC liposome which gave its permeability as 3.5×10^−13^cm^3^/s per single channel. The direct computation of the same authors produced 3.3×10^−13^cm^3^/s[37] at 25°C. These numbers compare well with our result of 7.67×10^−13^cm^3^/s (Table I and SI, Fig. S3) at the same temperature for AQP4 embedded in a POPE lipid bilayer. The difference between our results on AQP4 and the experimental data of Ref. [37] may have come from the fact POPE and POPC have different effects on AQP4. AQP4’s permeability was shown to be sensitive to the lengths of the POPC lipids used in Ref. [37]. The POPC head group is significantly bulkier than the POPE head group. We suggest that POPCs of longer tails have weaker stress on the protein’s extracellular and cytoplasmic ends and thus do not suppress AQP4 permeability as much as the shorter POPC lipids used in Ref. [37]. In contrast, POPE lipids do not exert a significant stress on the ends of AQP4. Correspondingly, this water channel embedded in a POPE bilayer has a higher permeability than the cases of POPC lipids.

In the many computational studies of the current literature including two studies[87, 88] using kilobar hydraulic pressure and many studies[17, 22, 50, 51, 57, 79, 89-91] using the theoretical formalism of single-file diffusion[92], the consensual results are around a fraction of 10^−13^cm^3^/s per single channel for AQPs 1, 4, 5, Z, and GlpF (when not occluded by glycerol). In Fig. 3 and SI, Figs. S6 to S8, we show the results of single-file diffusion from our large-scale simulations, which are all consistent with the current literature on smaller model systems (Table I). This agreement between the large and small systems indicates that the single-file diffusion approaches have reached convergence. Since the converged results are significantly away from the latest experimental data, we consider the possibility that water permeation through AQPs under osmotic gradient not be exactly single-file diffusion. We examined the water flows in the single-file channels and noticed constant fluctuations of the waters lining up approximately in single file inside a water pore by the hydrogen-bonding network with the pore-wall residues and with one another, which are definitely not rigid or in perfect order. Therefore, we conclude that the theoretical approximation of single-file diffusion gives us qualitative and fundamental understanding of aquaporin permeability but not quantitative accuracy. Direct computation of osmotic flux in sub M concentration gradient is necessary to produce quantitative evaluations of the aquaporin permeabilities.

**Fig. 3.**
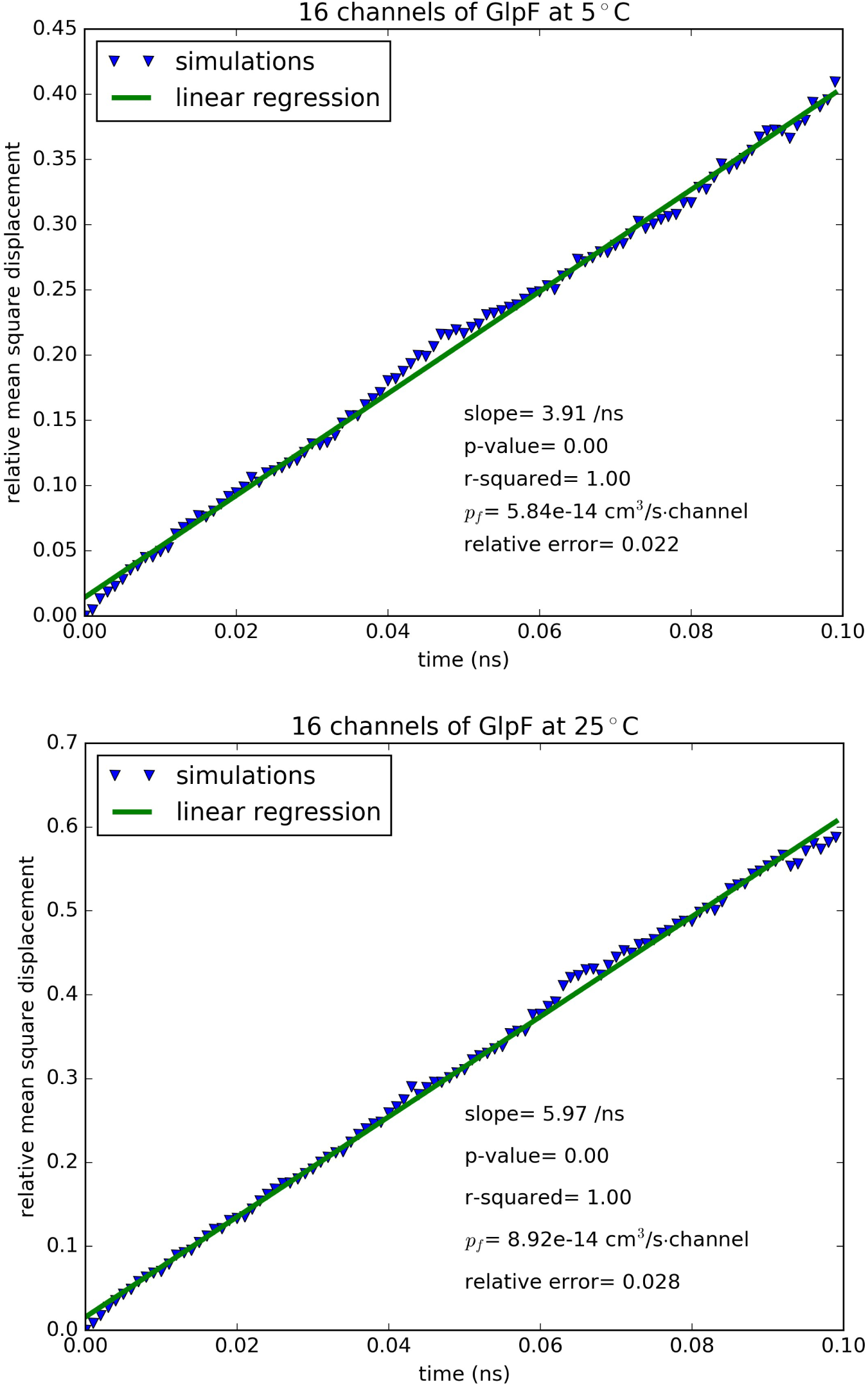
Mean square displacement of waters inside the single-file region of the GlpF channel: (A) Top panel, 5°C. (B) Bottom panel, 25°C.

**Second, the Arrhenius activation barriers** were experimentally determined to be around 3 kcal/mol for all water channels except AQP4 for which it is around 5 kcal/mol (Table II). The Gibbs free-energy computations all gave lower values in the free-energy barrier. This does not indicate that the free-energy computations were significantly inaccurate but means that the entropic contribution, TΔS (proportional to the absolute temperature T), is not negligibly small. The entropy barrier ΔS is opposite in sign to the energy barrier (the highest energy minus the lowest energy within the channel region). The entropy at the location of the highest energy is actually lower than the entropy at the location of the lowest energy. The experimentally measured Arrhenius activation barrier should not be compared with the free-energy barrier directly but with the activation energies extracted from the permeability computations. The agreements between our computed and the experimentally measured activation energies (Table II) are excellent in all three cases, which indicate the accuracy of the force fields in the current literature for water-protein interactions and the validity of our large-scale simulations.

### Third, what are the structural basis for AQP4’s strong temperature-sensitivity and high permeability?

In Refs. [70, 71], experiments already indicated that AQP4 has higher permeability than other water-specific channels. The experiments also showed that AQP4 has an Arrhenius activation barrier about 5 kcal/mol.[67, 71] A higher activation barrier means that AQP4 is much more temperature-sensitive than other water channel proteins. Namely, when the temperature is elevated, AQP4 increases its permeability more than other aquaporins. These characteristics of AQP4 correspond well its central role in keeping the hydrohomeostasis of the central nervous system, particularly at elevated temperatures. Our direct computation of permeabilities under near-physiological conditions is in perfect agreement with both of these AQP4 characteristics. Moreover, we computed the distances between the Arg and His residues that form the ar/R selectivity filter motifs of AQP4 and AQP5 (shown in Fig. 5). The significantly smaller fluctuations in AQP4 than in AQP5 well agree with the experimental data that the ar/R motif of AQP4 is more rigid than the ar/R motif of AQP5. The beta factors of AQP4-Arg216 are around 23[8] in contrast with AQP5-Arg188 having beta factors around 30[82].

**Fig. 4.**
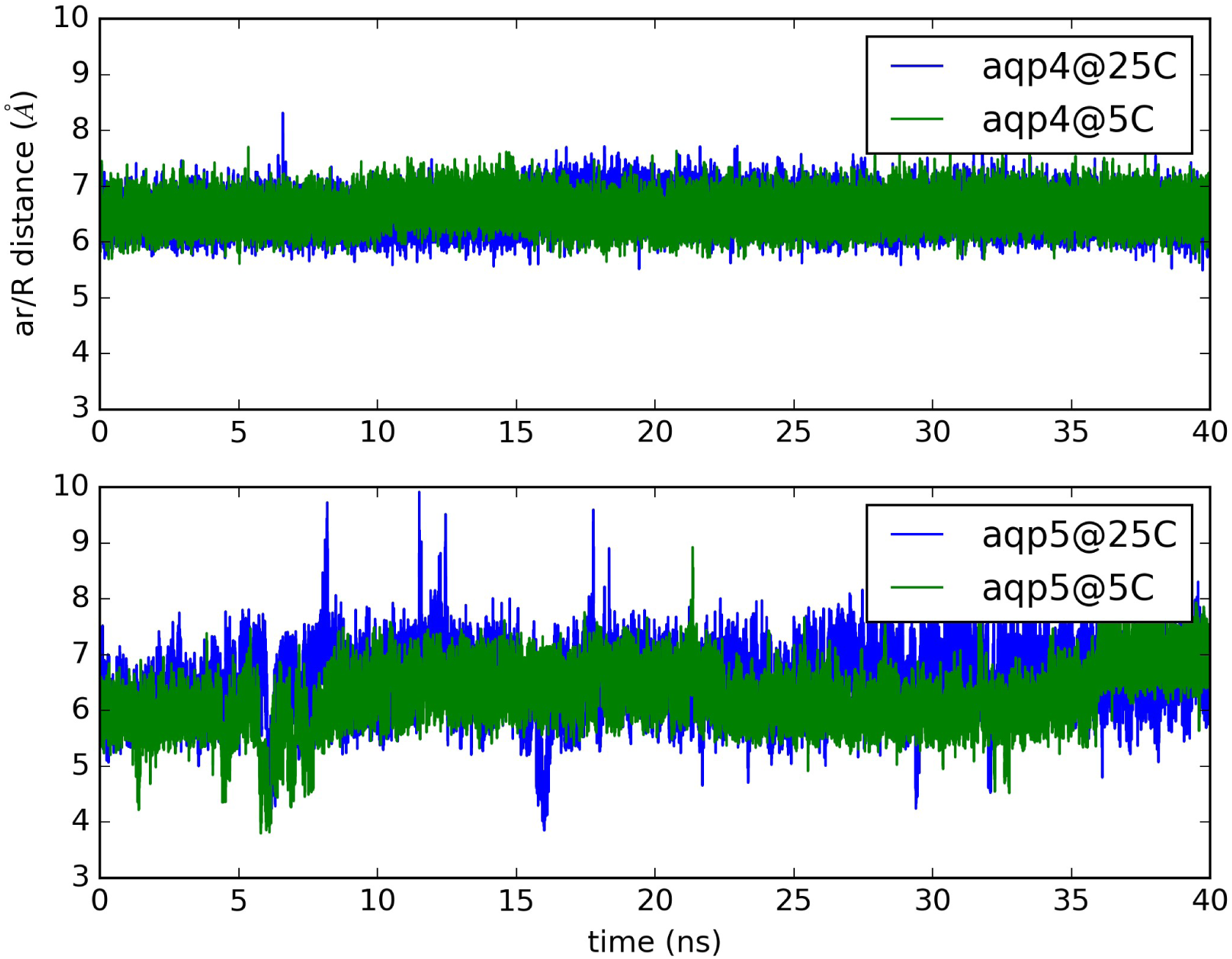
Fluctuations of the ar/R selectivity filter of AQP4 (top panel) and AQP5 (bottom panel). AQP4: distance between CZ of Arg 216 and CE1 of His 201. AQP5: distance between CZ of Arg 188 and CE1 of His 173.

## CONCLUSIONS

Through large scale simulations, we drew the following conclusions. In terms of methods, a model system of four aquaporin tetramers (16 channels) is large enough to have a signal to noise ratio for unambiguous results of osmotic currents induced by sub M concentration gradients across the membrane. The results of near-physiological simulations do not contradict but quantitatively differ from the far out-of-equilibrium simulations using kilo-bar hydraulic pressure. They also differ from the theoretical approximations on the basis of single-file diffusion inside the water channels. In terms of biophysical insights, AQP4, which plays a dominant role in hydrohomeostasis of the central nervous system is much more temperature-sensitive and more permeable than other water-specific channel proteins, *e.g.,* AQP5 that is largely responsible for water transport in saliva glands. Interestingly, AQP4 and AQP5 are structurally very similar in terms of hydrogen-bond networks formed by the waters inside a conducting pore with the pore-lining residues. Yet, they have very different temperature sensitivities due to differing dynamics behaviors of the ar/R motif residues. Indeed, minor differences in the structures-dynamics of a protein can cause significant differences in its biological functions.

## SUPPLEMENTAL INFORMATION

Additional figures are shown in Figs. S1-S13 of the SI for AQP4/5 system setup, simulation data, AQP4’s free-energy profile, and the differences between AQP4 and AQP5 in their ar/R dynamics (fluctuations).

## ACKNOWLEDGEMENTS

The authors acknowledge support from the NIH (GM 084834 and GM 060655) and the computing resources provided by the Texas Advanced Computing Center at University of Texas at Austin.

